# Visualization of translation reorganization upon persistent collision stress in mammalian cells

**DOI:** 10.1101/2023.03.23.533914

**Authors:** Juliette Fedry, Joana Silva, Mihajlo Vanevic, Stanley Fronik, Yves Mechulam, Emmanuelle Schmitt, Amédée des Georges, William Faller, Friedrich Förster

## Abstract

Aberrantly slow mRNA translation leads to ribosome stalling and subsequent collision with the trailing neighbor. Ribosome collisions have recently been shown to act as stress sensors in the cell, with the ability to trigger stress responses balancing survival and apoptotic cell-fate decisions depending on the stress level. However, we lack a molecular understanding of the reorganization of translation processes over time in mammalian cells exposed to an unresolved collision stress. Here we visualize the effect of a persistent collision stress on translation using *in situ* cryo electron tomography. We observe that low dose anisomycin collision stress leads to the stabilization of Z-site bound tRNA on elongating 80S ribosomes, as well as to the accumulation of an off-pathway 80S complex possibly resulting from collision splitting events. We visualize collided disomes *in situ*, occurring on compressed polysomes and revealing a stabilized geometry involving the Z-tRNA and L1 stalk on the stalled ribosome, and eEF2 bound to its collided rotated-2 neighbor. In addition, non-functional post-splitting 60S complexes accumulate in the stressed cells, indicating a limiting Ribosome associated Quality Control clearing rate. Finally, we observe the apparition of tRNA-bound aberrant 40S complexes shifting with the stress timepoint, suggesting a succession of different initiation inhibition mechanisms over time. Altogether, our work visualizes the changes of translation complexes under persistent collision stress in mammalian cells, indicating how perturbations in initiation, elongation and quality control processes contribute to an overall reduced protein synthesis.

**Summary:** Using *in situ* cryo electron tomography we visualized the reorganization of mammalian translation processes during a persistent collision stress.

## Main

In mammalian cells, several ribosomes typically translate the same mRNA molecule in an arrangement called polysome. In situations like a defect in mRNA, a repetition of slow codons, or nutrient starvation, the ribosome stalls on the mRNA which induces a collision with its trailing neighbor (Fig. 1A). Ribosome collisions have recently been shown to act as sentinel for stress situations by triggering general stress responses affecting cell fate (*1, 2*) (Fig. 1A). Ribosome collisions occur under physiological growth conditions (*3, 4*), triggering quality control mechanisms for their clearing. The Ribosome associated Quality Control (RQC) pathway detects the truncated nascent chain resulting from stalled translation and ubiquitinates it for proteasomal degradation. More recently, different intensities of ribosome collision stress were shown to induce distinct general stress responses (*1, 5*) (summarized in Fig. 1A). A low intensity collision stress triggers the Integrated Stress Response (ISR) for survival, resulting in the phosphorylation of eIF2α which is known to efficiently inhibit translation (*6*). Upon a stronger persistent collision stress, the Ribotoxic Stress Response is activated, resulting in the phosphorylation of p38 and JNK, leading to cell cycle arrest and apoptosis (*1, 5*).

**Figure 1:**
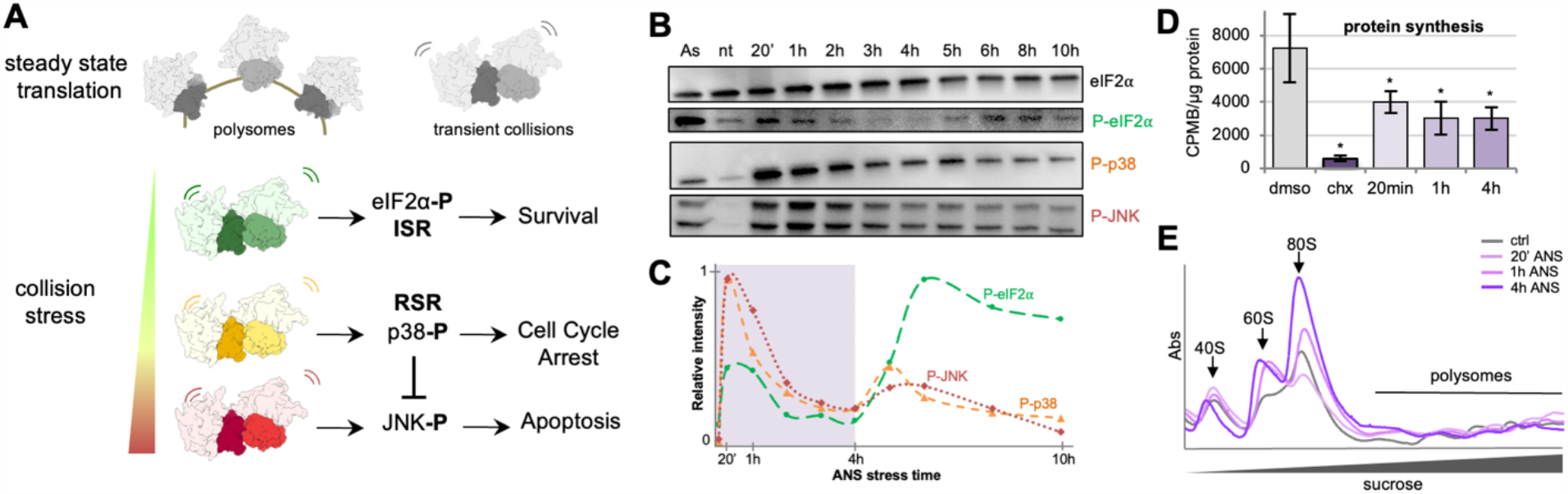
Biochemical analysis of low dose ANS persistent collision stress in MEF cells. (A) Schematic representation of translational situation in untreated cells (polysomes and low amount of collisions) and under increasing intensities of collision stress with associated cellular stress responses and cell fate outcome. (B) Western blot analysis of collision stress induced responses in MEF cells treated with 500 µM Arsenite for 20 min, untreated, or treated with 200 nM ANS for 20 min to 10h: total eIF2α, phosphorylated eIF2α, phosphorylated p38 and phosphorylated JNK (C) Corresponding relative intensity measurements, background substracted and normalized by total eiF2α intensity (D) ^35^S incorporation protein synthesis measurements in control cells and cells treated with high dose CHX (100 mg/mL), and low dose ANS (200 nM) for 20min, 1h and 4h. (E) Polysome profiling on sucrose gradients for control cells and cells treated with 20min, 1h and 4h low dose ANS.

The molecular factors involved in ribosome collision detection and the subsequent triggering of general stress responses have also been a recent topic of investigations. In mammalian cells, the abundant EDF1 binds to early collisions at the mRNA entry channel on the 40S subunit near the collision interface and recruits the translation repression factors GIGYF2 and eIF4E2, thereby preventing further initiation on the problematic mRNA (*7, 8*). GCN1 further binds collided disomes and acts as a checkpoint, triggering the activation of GCN2 in the case of persistent collisions (*9*). The long isoform of the MAPKKK ZAKα recognizes stalled ribosomes and is required for the activation of both the ISR and the RSR (*1, 10*). Next, the E3 ligase ZNF598 ubiquitinates RPS10 at the collision interface of persistent disomes (*3, 11-13*). RPS10 ubiquitination is a signal recognized by the ASC-1 Complex (ASCC) (*14-16*) which pulls on the mRNA and splits the stalled ribosome by pushing it against its collided neighbor (*17*). The soluble factor NEMF then recognizes the stalled 60S complex and recruits the low abundance E3 ligase listerin for ubiquitination of the truncated nascent chain (*18, 19*), finally targeted to proteasomal degradation. This recently elucidated network of responses suggests a subsequent reorganization of the translation processes in a cell under persistent collision stress, which has not been explored so far. For instance, it is unclear how much collisions can accumulate in the cell, to which extent this accumulation is limited by feedback mechanisms and for how long the cell can survive a persistent collision stress. Using *in situ* cryo electron tomography we visualized in mammalian cells the evolution of the ribosomal populations over time during a low dose anisomycin (ANS) stress, providing a molecular basis for the translational reorganization under persistent collision stress.

### Low dose ANS collision stress induces dynamic signaling of ISR and RSR in MEF cells

To characterize persistent collision stress in mammalian cells, we used MEF cells treated with low dose ANS (200nM ANS) for 20 min to 10h. We observed no sign of cell death under these conditions (no cell detaching from the plate, no cleavage of apoptotic markers PARP and Caspase 3, Fig S1A). We analyzed the activation of the ISR and RSR pathways over time by Western blotting against corresponding markers (Fig. 1B). These experiments reveal 2 distinct peaks of the stress response (Fig. 1B, 1C). The first ISR and RSR responses occur early, as indicated by peaks in the phosphorylation levels of eIF2α, p38 and JNK at 20 min stress. However, when the collision stress persists, these phosphorylation levels go down over time, with P-eIF2α reaching a minimum at about 4h stress, followed by a second ISR wave. We hypothesized that cellular survival mechanisms might be initiated during the first stress response, and the failure to re-establish homeostasis within ∼4h may trigger the second stress response. We focused our analysis on the progression of the translational situation in cells exposed to low dose ANS stress between the previously studied early collision stress timepoint of ∼20 min (*1*) and the 4h turning point. We measured total protein synthesis rates under these stress conditions following ^35^S-methionine incorporation (Fig. 1D). As a control for the stalling of most ribosomes in the cell, we used a high dose (100 μ g/mL) of cycloheximide (CHX), which decreased protein synthesis about 12-fold when compared to DMSO treated control (p=0.015). Low dose ANS stress at all timepoints led to a more modest 2-fold decrease of protein synthesis (15min p=0.049, 1h p=0.03, 4h p=0.03), indicating that translation initiation and/or elongation rates are decreased. To further analyze the effect of low dose ANS stress on translation in mammalian cells, we performed polysome profiling on control MEF cells and cells submitted to 20 min, 1h, 4h low dose ANS stress, revealing an accumulation of isolated 80S and 60S in stressed cells over time (Fig. 1E). Western blot analysis of the corresponding polysome fractions suggested an increase of RPS10 ubiquitination on polysomes under stress (Fig. S1B), consistent with previous studies of collisions stress (*3, 12*). In addition, we detect an accumulation of EDF1 (but not ZNF598) on stressed polysomes over time (Fig. S1C), in good agreement with previous observations (*7, 8*).

Altogether our data suggest that MEF cells reorganize their translation over time under persistent low dose ANS collision stress.

### *In situ* cryo electron tomography reveals the use of the Z-site during elongation in MEF cells

To first analyze the baseline ribosomal elongation situation in control MEF cells, we used *in situ* cryo-electron tomography. We obtained 28,644 ribosomal particles which altogether yielded a subtomogram average of the ribosome at ∼6.7 Å resolution (Fig. 2A, 2B). To increase the number of particles and obtain a resolution benchmark for our setup, we pooled these control particles with the 4h ANS dataset (18,285 particles), leading to an improved resolution down to ∼4.5 Å in the best resolved part of the LSU (Fig. S2, S3). We next used classification to separate the distinct SSU rotation and tRNA occupation states of the 80S ribosomes from untreated cells. The resulting 9 main classes display clear neighboring ribosome densities indicating that they can be found on polysomes and correspond to eukaryotic 80S elongating complexes as previously reported (*20-22*) (Fig. 2C, 2D, 2E, S4): a decoding E state (16035 particles, 8.8 Å resolution) (*20, 23, 24*), a classical PRE state with or without a less well-defined extended elongation factor density (130 to 2183 particles per class, resolution 26 Å to 9.2 Å, Fig. S5). (*20, 21, 23, 25, 26*), a rotated-2 state bound by eEF2 (2,184 particles, 9.8 Å, rotated-2 +) or without (409 particles, 15.1 Å, rotated 2) (*20-22, 26*), an unrotated translocation intermediate (663 particles, 13.9 Å, POSTi) (*20, 21*), and a low abundance unrotated POST state (178 particles, 19.7 Å) (*20, 21*).

**Figure 2:**
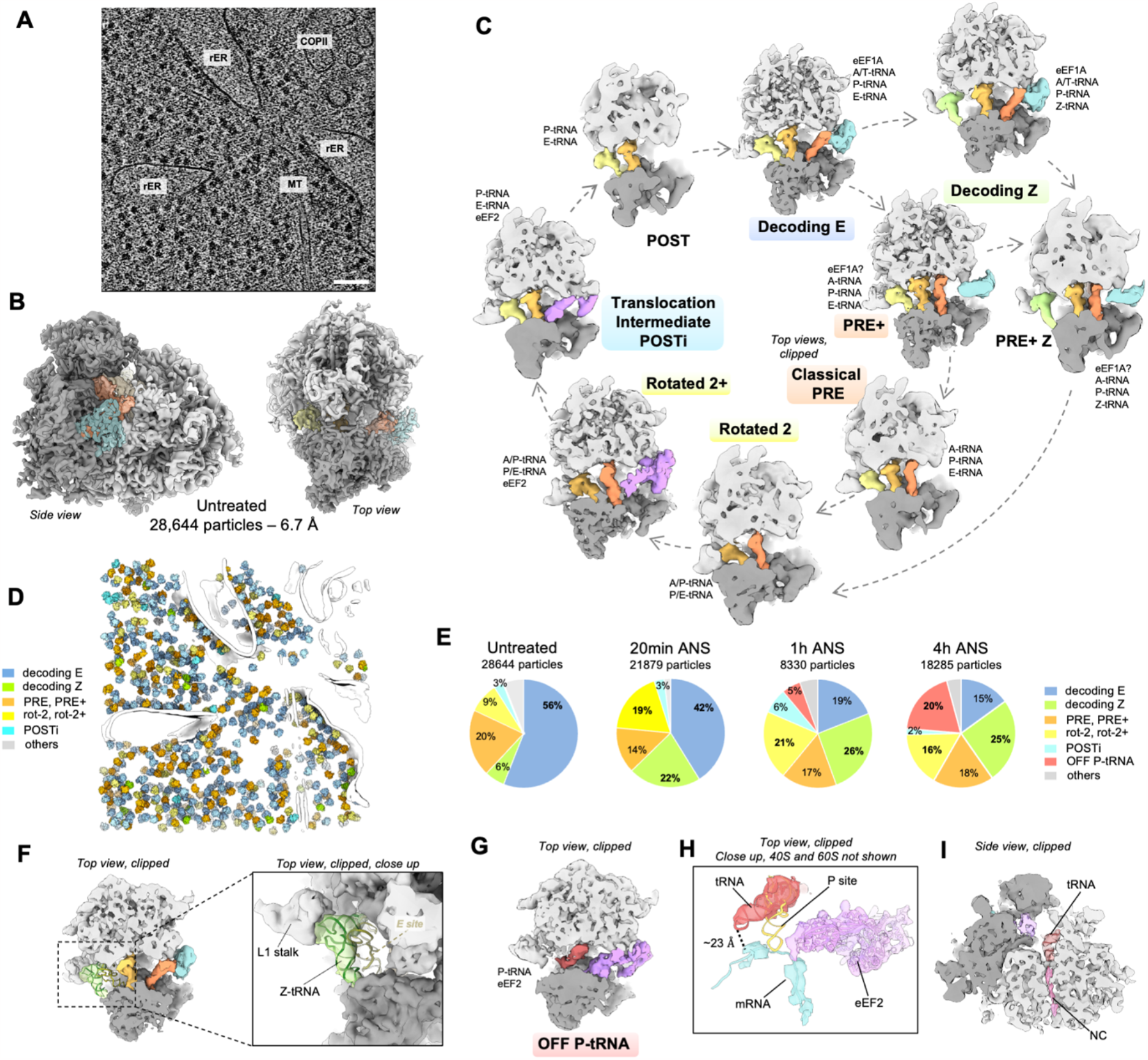
*In situ* visualization of 80S ribosome populations. (A) Slice through a representative tomogram of control MEF cells, scale bar: 100 nm. (B) Subtomogram average of 80S particles in the control condition. The small subunit is displayed in dark gray, the large subunit in light gray, the elongation factor in cyan and the tRNAs in shades of orange to yellow. (C) Observed active intermediates positioned in model of mammalian elongation cycle. The ribosome is clipped for visualization. A, P, and E indicate ribosomal aminoacyl, peptidyl, and exit sites, respectively. The tRNAs are color-coded with respect to a complete cycle. The color code is the same as in B, with eEF1A in cyan, eEF2 in purple and the Z-tRNA in green. (D) Different ribosome elongation states mapped back in the original tomogram shown in A. Segmented membranes and microtubules are displayed in white. Relative abundance of ribosomal elongation complexes in all datasets. (F) Close-up view on the Z-site bound tRNA of the decoding Z complex. (G) Off-pathway ribosomal complex observed under prolonged low dose ANS stress (1h and 4h). The tRNA is displayed in dark red. (H) Same complex as in G, displaying a model fit for the tRNA, eEF2 and the mRNA density (dark cyan). (I) Same complex as in G and H, side view, clipped for the visualization of the peptide exit tunnel displaying a nascent chain (pink) bound to the tRNA.

In addition, we observe an 80S complex present on polysomes and similar to a decoding state, with the only difference that no tRNA is present in the E-site. Instead a tRNA is bound to a site further out and in contact with the ribosome head and L1 stalk, previously described as Z-site (1,722 particles, 10.3Å, decoding Z, Fig. 2B, 2F, S4) (*27*). We also observe a smaller PRE+ like class carrying a Z-site tRNA (382 particles, 15.4 Å, PRE+ Z) and with enriched density for trailing neighbor when compared to leading one, suggesting that this low abundance class may be enriched in transiently stalled ribosomes. Due to the similarity of interactions between the Z-site and the binding site of mRNA Internal Ribosome Entry Sites (IRES), it was proposed that the Z-site is a preferred binding sites for certain RNA structures possibly including cellular mRNAs (*27*). Alternatively, it could also be the result of de-acylated tRNA re-binding (*27*). So far, the Z-site had only been observed in ribosomes purified from rabbit reticulocyte lysates, limiting the functional relevance of these observations. Our results show that the Z-site is a functional tRNA site occupied *in situ* in ribosomes present on polysomes and with a P-site tRNA, under normal growth conditions in MEF cells.

### Low dose ANS stress is associated with increased abundances of Z-site bound states and apparition of a monosomal 80S complex

To investigate the effect of low anisomycin stress on native ribosomal populations we performed the same analysis on datasets collected on ANS-stressed MEF cells (Fig. 2E, S3). We focused our analysis on the 4h timepoint where cells persist despite a first wave of ISR and RSR (Fig. 1B, C, E) and our polysome profiling suggested an increase in isolated 80S. We compared this 4h stress timepoint with the 20min and 1h datasets used as indicative of earlier intermediate stress timepoints. All main populations described for the control cells were also found in polysomes in stressed cells. However, their relative abundance is affected by the ANS stress over time, indicating possible kinetics barriers introduced by the elongation stress (Fig. 2E).

We observe an increase in the rotated populations (from 9% to >16%) in stressed cells when compared to normal growth conditions. This is consistent with the possible accumulation of ribosome collisions, where the collided ribosomes are found in the rotated-2 conformation (*3, 9, 12, 28-30*). The decoding Z state abundance is also increased under stress (from 6% to about 25% of the total 80S populations, 4h ANS: 3,461 particles 8.9 Å). An overall decrease of translation elongation rate in the cell could favor the stabilization or re-binding of a deacylated tRNA in such an intermediate exit position. Alternatively, mRNAs expressed as a response to stress may preferentially use the Z-site for their translation. We also observe that at 4h ANS stress, classical PRE+ states are found with a Z-site bound tRNA or displaying empty Z- and E-sites. In addition, the 4h ANS PRE+ and PRE classes display a weak leading neighbor density but a strong trailing one, indicating that these classes are enriched in stalled ribosomes (Fig. S4).

Finally, a previously undescribed 80S conformation accumulates at long stress timepoints (∼5% after 1h and ∼20% at 4h, 3573 particles at 4h, 9 Å, Fig. 2G). This class displays a tRNA bound to the 60S P-site but no density for polysome neighbors (Fig. S4), indicating that it is not actively elongating. The presence of this abundant monosomal 80S complex is likely to account for the increase in 80S peak we observed by polysome profiling at 4h stress (Fig. 1E). This class further displays partial density for mRNA, but the anticodon loop of the P-tRNA is not interacting with the mRNA, instead pointing away from it (∼23 Å distance, Fig. 2H). The tRNA acceptor stem bound to the 60S P-site is attached to a nascent chain visible in the 60S peptide tunnel (Fig. 2I), indicating previous elongation activity. We speculate that these ribosomes might be the result of ASCC splitting of previously collided disomes. The pulling force of the ASCC on the mRNA to split the stalled ribosome, may have detached the mRNA from the A/P tRNA in the second collided ribosome (*17*). Further studies will be required to investigate the cellular impact of the accumulation of these ribosomes and their possible elimination mechanisms such as autophagic recycling (*31*).

### Ribosome collisions occur upon polysome compression and reveal an interface between the Z-site tRNA and L1 stalk on the stalled ribosome and eEF2 on the rotated-2+ collided neighbor

To analyze the spatial distribution of ribosomes in cells and its changes under low dose ANS stress, we performed nearest neighbor analysis. We calculated distances between an 80S mRNA entry site and the closest exit site on other 80S in the proximity and the other way around. We selected the shortest distance for counts and used a 12nm cutoff to focus our analysis on distances compatible with neighbor particles on the same polysome (∼7 nm). While most of the distances in the control dataset are spread across a wide peak centered around ∼7 nm (Fig. 3A), we observed that low dose ANS stress induces a continuous compression of these distances over time (Fig. 3A-D). The corresponding plots display a narrow peak at ∼4.5 nm distance after 1h and 4h stress, compatible with mRNA length from entry to exit sites on collided disomes (*3, 12, 30*).

**Figure 3:**
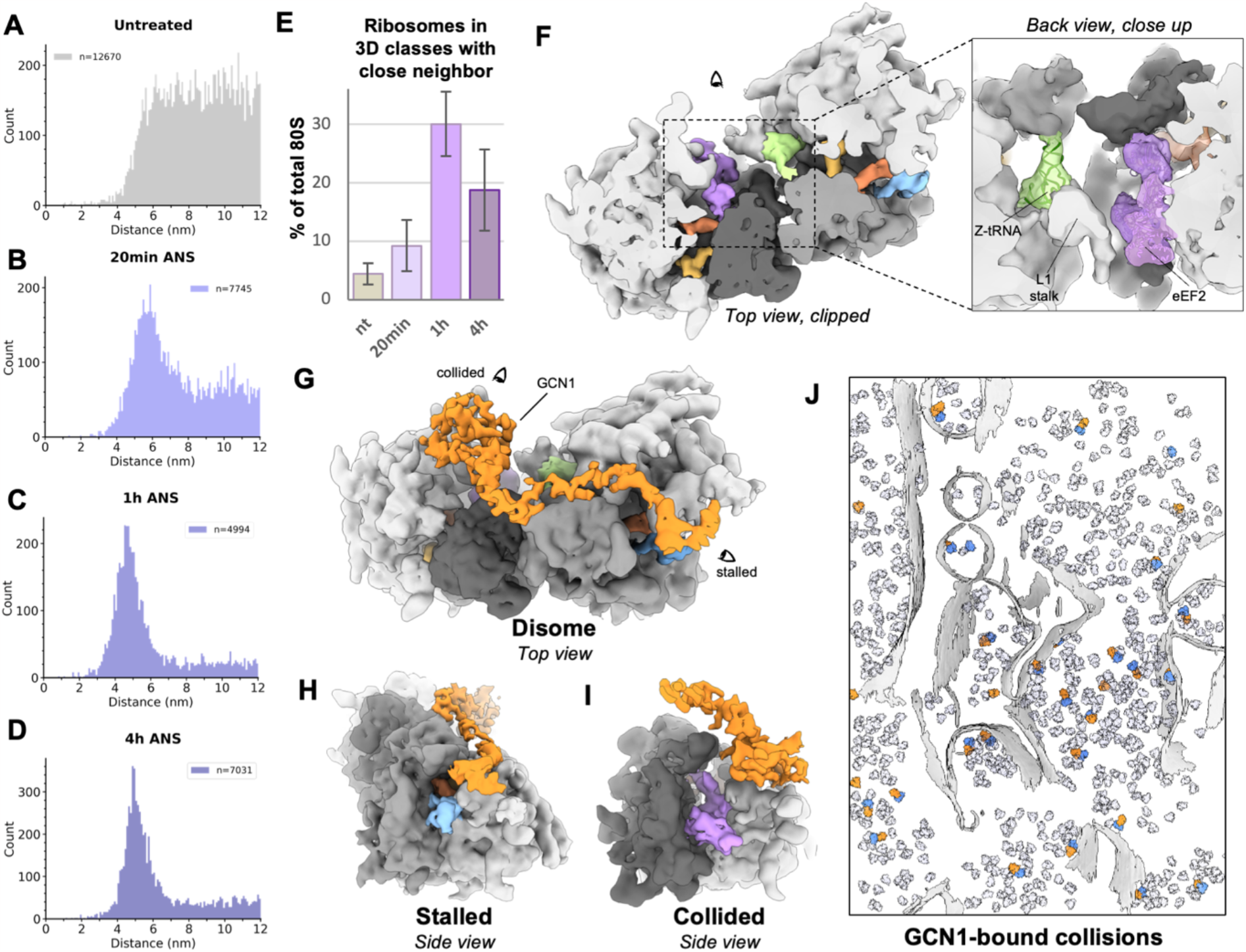
*In situ* analysis of ribosome collisions. (A) Distance to nearest neighbor plot for the control data set, (B) same at 20 min low dose ANS stress, (C) same at 1h low dose ANS stress, (D) same at 4h low dose ANS stress. On each plot n indicates the number of distances counted in the plot, ie. all entry/exit distances < 12 nm. (E) Rough quantification of ribosomes with a defined close neighbor based on RELION 3D classification results. Bar and whiskers are mean and s.d. across tomograms (untreated n = 87, 20 min ANS n = 68, 1h ANS n = 36, 4h ANS n = 45). (F) *In situ* subtomogram average of collided disome. Close up view displays fitted models for the Z-site tRNA and eEF2. (G) *In situ* subtomogram average of GCN1-bound collided disomes and side views: (H) of the stalled ribosome, (I) on the collided ribosome. (J) GCN1-bound collisions mapped back into a tomogram. Segmented membranes are displayed in light gray and all other 80S particles in transparent light mauve.

To visualize the ribosome populations with close neighbors, we used image classification with a mask on the positions for neighboring SSUs (Fig. S6). We detected ribosomes with defined densities for contacting neighbors in all datasets, including the non-stressed cells, albeit in different amounts (Fig. 3E, S6-8): about ∼4.5% of 80S ribosomes in untreated cells – in good agreement with previous observations in vertebrate cells (*3, 8, 32*), followed by a ∼2-fold increase to ∼8.5% at 20min stress, a peak at ∼28% at 1h stress followed by a decrease to ∼16% at 4h stress. These counts are given as indicative, as our image classification is limited by the presence of some false positive and negative in the collision classes precluding accurate quantification (ribosomes from these classes plotted back into their original tomograms do not fully result in perfect pairing as seen in Fig. S6-7). The decrease in collisions at 4h stress may be the results of negative feedback mechanisms reducing protein synthesis (Fig. 1D) (*7, 8*).

To assess the elongation states of the ribosomes in disomes, we further used image classification with a mask around tRNA sites and GTPase Activation Center (GAC). Our analysis reveals that collisions visualized in low ANS treated cells are more diverse than anticipated from previous work on purified disomes (*30*). We distinguish two classes of stalled ribosomes: a major decoding-like state and a PRE+ like state. We also separate two classes of collided ribosomes: a major rotated-2+ state and another unrotated class, mostly populated by unrotated decoding like ribosomes (Fig. S6).

The most abundant disome arrangement in our data displays an overall geometry that is in very good agreement with the recently reported cryoEM structure of *XBP1u* stalled human disomes (Fig. 3F) (*30*). However, the stalled ribosome population in our data is enriched in a decoding-like state and displays a tRNA bound to the Z-site. The density for the factor bound to the stalled ribosome GAC is different from the typical compact eEF1A bound to previously described decoding states (Fig. 3F). This density is likely a factor involved in the recognition of stalled ribosomes, like DRG2 (Rbg2 in yeast) (*9*), but the limited resolution of our subtomogram average precludes its identification. On the collided ribosome, we observe the binding of eEF2 – which may have detached in previous purifications of mammalian collision complexes (*3, 30*). This preferred collision geometry appears to be stabilized by a contact formed by the Z-site bound tRNA and L1 stalk of the leading ribosome with eEF2 bound to the collided rotated-2+ neighbor (Fig. 3F). This interaction provides a possible explanation for the observed increase of the decoding Z class under low ANS stress (Fig. 2C, 2F) and for the apparent incapacity of the collided ribosome to complete the translocation step (*30*).

Our approach further visualized mammalian GCN1-bound collisions *in situ* which are similar to purified GCN1 bound yeast disomes (Fig. 3G-J) (*9*). GCN1-bound collisions represent ∼50% of total collisions in untreated cells, ∼30% at 20min stress, ∼20% at 1h and ∼30% at 4h stress. Hence, GCN1 bound disomes appear to be relatively long lived in cells, including in the absence of stress. Together with the changes in the proportion of collided ribosomes, these abundances suggests that GCN1 may also become limiting in detecting and clearing accumulating collided ribosomes.

In addition to the previously described disomes, we also observe a class of middle ribosomes displaying contacting neighbors in both the leading and the trailing positions (Fig. S6-8). This class corresponds to a compact helical polysomal arrangement (Fig. S7) (*33, 34*). These ribosomes are mostly found in a POSTi translocation intermediate step, indicating a possible slower translocation in this arrangement. The trailing and leading neighboring ribosomes appear at a similar distance, but less defined and with an angle distinct from the one observed in typical collided disomes (Fig. S7). We detect this class at 20min, 1h and 4h of stress (Fig. S6), but we could not isolate it in the non-stressed cells, where the ribosomes are more loosely packed (Fig. 3A) and classes with visible close neighbors are in lower abundance (Fig. 3E). Modelling longer chains of ribosomes show that the typical collided disome arrangement can only accommodate up to 4 ribosomes without introducing severe clashes. On the contrary the helical geometry is not limited in polysome length (Fig. S7, S8). Our modelling further reveals that this helical arrangement is not compatible with GCN1 binding on top of the P-stalk of the trailing ribosome (Fig. S7). Hence, our data suggests that the helical polysome arrangement can accommodate closely packed polysomes, for instance to buffer situations like the slow-down of a leading ribosome, without triggering GCN1 induced stress responses.

Finally, we visualized ER-bound collisions, which appear in good agreement with the previously proposed hypothesis that collided pairs of ER-bound ribosomes adopt the same geometry as cytosolic disomes, relying on a local negative curvature of the fluid ER membrane (*3*) (Fig. S9).

### Non-functional t-RNA bound 60S particles accumulate under persistent collision stress

Polysome profiling in MEF cells revealed the accumulation of free 60S under persistent collision stress (Fig. 1E). To investigate the possible reasons for their accumulation we analyzed 60S particles in our *in situ* cryo-ET datasets. A similar pattern was observed at all stress timepoints (20 min, 1h, 4h) and we focused our detailed analysis on the 4h dataset. We detected about 1500 and 2700 60S particles, respectively in the control and the 4h low ANS stress datasets. In the untreated cells, the major population detected (∼64%) corresponds to a complex of the 60S particle bound by a density interacting with RPL23 displaying a shape and position compatible with eIF6, as well as an extra factor at the GAC, and a density probably corresponding to EBP1 at the exit tunnel (Fig. 4A, 4C). It is likely that this complex represents a late cytosolic maturation state of the large ribosomal subunit, possibly bound by SBDS and EFL1 for the release of eIF6 and the formation of a 60S particle fully competent for subunit joining. In addition, we detected another population of 60S strongly resembling the earlier cytosolic maturation intermediate state B (∼10%, Fig. 4B, 4C) (*35*), displaying densities matching LSG1, NMD3, eIF6, ZN622 and EBP1. After 4h of low ANS stress, the proportion of state B maturation intermediate is decreased, and the idle 60S later maturation intermediate is not observed anymore. Instead, 60S particles appear bound by a P-site tRNA, a protein nascent chain, eIF6 and EBP1 densities (Fig. 4D, 4F, S10), likely resulting from collision splitting reactions. To analyze the RQC response to the collision stress, we further classified the 60S populations of each dataset. We identified a sub class displaying densities typical for NEMF and listerin in size, shape and location (*19*) and accounting for ∼10% of the total 60S populations in both the untreated and 4h ANS datasets. In the 4h ANS dataset, we detected additional tRNA bound populations possibly also bound by NEMF alone (Fig. S10). These observations confirm that under persistent collision stress, listerin is limiting in resolving the post-splitting 60S complexes containing a tRNA and a nascent chain (*18, 19*), leading to the accumulation of these non-functional 60S complexes in the cytosol. In addition, our data indicates that under persistent collision stress, late LSU maturation intermediates, generating subunit joining competent 60S complexes upon eIF6 release, may become limiting in the cell, thereby possibly contributing to the observed overall decrease in protein synthesis (Fig. 1D). Further studies will be required to investigate how this pool is regulated under collision stress.

**Figure 4:**
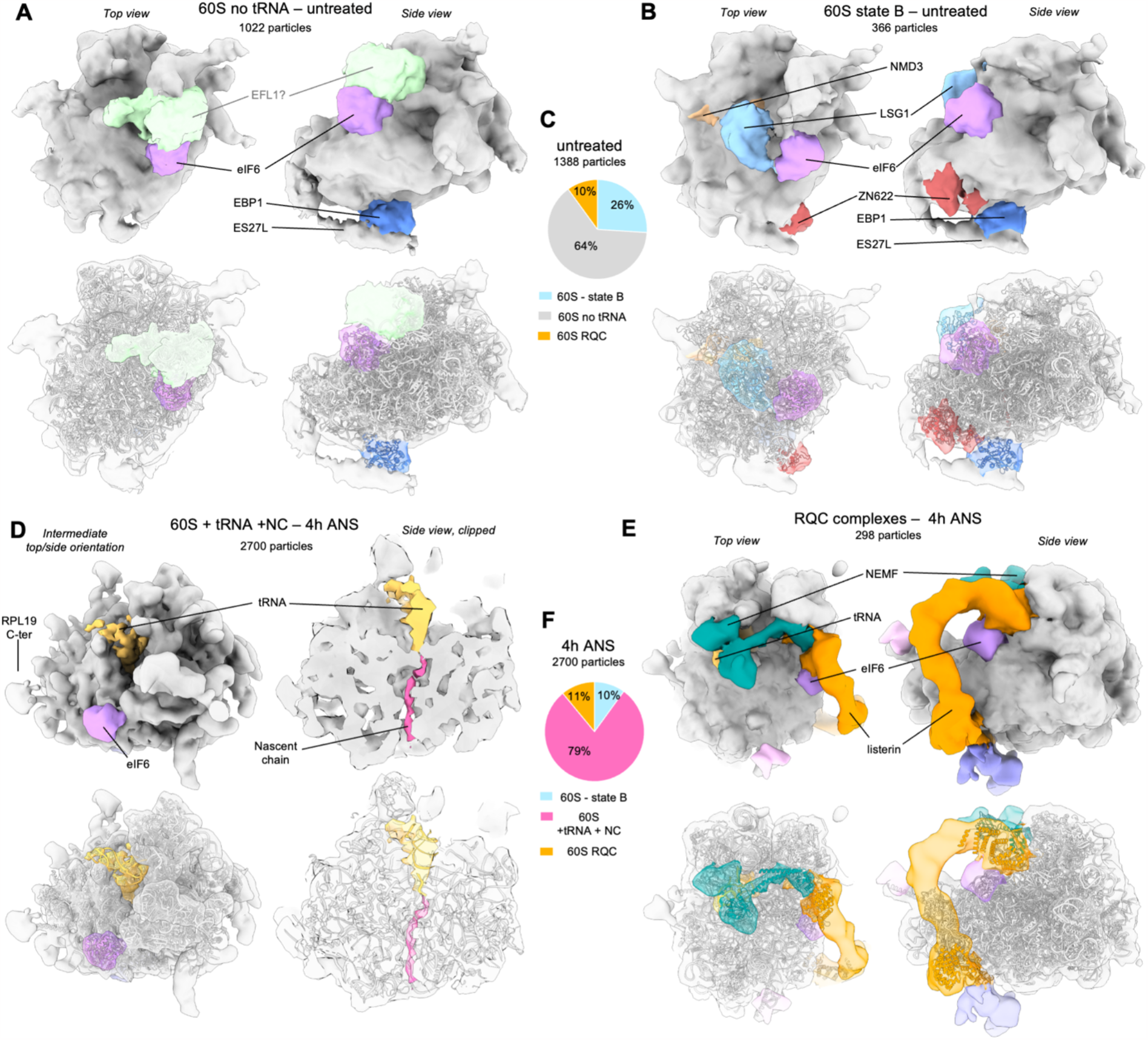
*In situ* subtomogram averages of 60S complexes. (A) Major 60S complex observed in control cells displaying densities corresponding to eIF6 (purple), EBP1 (blue) and a putative eFL1 (light green). PDB coordinates for 60S, eIF6 and EBP1 were fitted independently (using the corresponding chains from PDB 7OW7 for the 60S and eIF6, and 6LSR for EBP1). (B) Second most abundant 60S class, corresponding to previously described maturation state B, displaying densities corresponding to eIF6 (purple), LSG1 (light blue), NMD3 (beige), ZN622 (dark red). Fitted PDB coordinates: 6LSR. (C) Relative abundances of the 60S complexes observed in the control dataset. (D) Major 60S complex observed in the 4h low dose ANS stress dataset displaying densities for eIF6 (purple), partial P-site tRNA (gold) and nascent chain (hot pink). PDB coordinates were fitted independently using the corresponding chains from 3J92 for 60S, eIF6 and tRNA, and 5AJ0 for the nascent chain. (E) NEMF and Listerin bound 60S particles as observed in stressed cells. Fitted PDB coordinates: 3J92. (F) Relative abundances of the 60S complexes observed in 4h low dose ANS stress dataset.

### Low dose ANS stress induces different perturbations of translation initiation over time

Cell stress has also been shown to impacts translation initiation, including via the ISR. Phosphorylation of eIF2α induces the formation of a stable inactive complex between phosphorylated eIF2-GDP and the recycling factor eIF2B present in limiting amounts, inhibiting canonical initiation and contributing to a decrease in protein synthesis (*36-38*). We used subtomogram averaging and classification to analyze possible molecular differences between the small subunit complexes in untreated cells and under low dose ANS collision stress. In all datasets, we detected 40S and 43S complexes (Fig. 5A-F). As these populations are much less abundant than 80S populations, quantifications are indicative, and we focused our analysis on the main differences between normal and stress conditions. While the proportion of 40S complexes remain relatively similar across conditions (Fig. 1E), we note apparition of new classes of particles under stress, with important shifts in the proportion of those complexes between timepoints.

**Figure 5:**
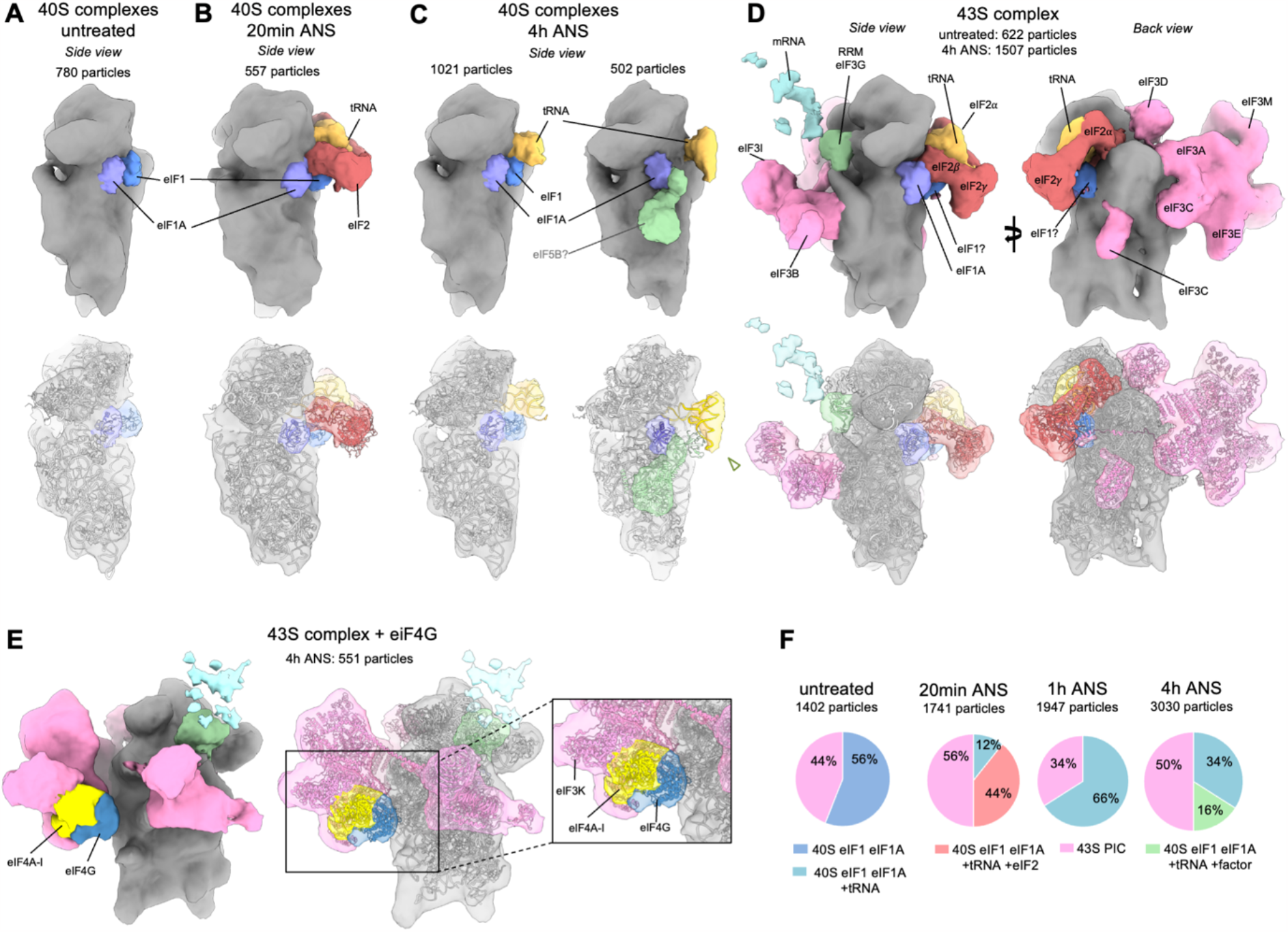
*In situ* subtomogram averages of 40S complexes. (A) 40S complex observed in control cells, densities corresponding to eIF1 and eIF1A are displayed in blue and purple respectively. Fitted PDB coordinates: 4KZY. (B) Abundant 40S complex appearing at 20 min low dose ANS stress, displaying extra densities corresponding to tRNA (gold) and eIF2 (red). (C) Major 40S complexes observed at 4h low dose ANS stress displaying a P-site tRNA (gold) and a density depicted in light green, possibly fitting eIF5B (PDB coordinates: 7TQL). (D) 43S complexes observed in all datasets, displaying eIF3 (pink), and an extra density shown in light cyan proposed to correspond to an mRNA. Fitted PDB coordinates: 6ZMW. (E) Subtomogram average of a 4h ANS 43S subclass. The density corresponding to eIF4A1 is displayed in yellow and eIF4G in dark blue. Fitted PDB coordinates: 6ZMW. Relative abundances of the 40S complexes observed in untreated cells and under low dose ANS stress timepoints of 20min, 1h and 4h.

In the control cells, the largest population we observe is a 40S subunit which seems bound to eIF1 and eIF1A (Fig. 5A).

At 20 min low dose ANS stress, we observed the apparition of a 40S complex with densities resembling: eIF1, eIF1A as well as eIF2 (but not eIF3) and a P-site tRNA (Fig. 5B). This complex is only observed in our 20 min stress dataset, when a fraction of eIF2α is phosphorylated (Fig. 1B). The affinity of eIF2-GDP for the methionine tRNA was shown to be 150nM vs. 9nM for eIF2-GTP (*39*). We may thus be observing eIF2-GDP bound in this 40S complex, which could hamper the following steps of initiation until eIF2α phosphorylation level decreases (Fig. 1B).

Later at 4h ANS, after the first phosphorylation peak of eIF2α, our dataset displays 2 distinct populations of 40S complexes (Fig. 5C). The first one seems bound by eIF1, eIF1A and a tRNA. The second one carries a tRNA in a different position and an additional factor. Structural comparison rather exclude that this factor could be ABCE1 or TSR1, while eIF5B and a de-acylated tRNA bound to this complex appear more likely (Fig. S11, 5C) (*40, 41*).

The major 43S complex observed is similar in all datasets, displaying densities resembling eIF1, eIF1A, eIF2α, eIF2β, eIF2γ, eIF3, eIF4A1 and a P site t-RNA (Fig. 5D) (*42-45*). In the untreated sample, we observe a strong density at the 40S mRNA entry channel. It starts around the position of an RNA Recognition Motif (RRM) and apparently interacts with eIF3g, eIF3i, possibly also eIF3b. eIF3 is required for scanning and in yeast where eIF3g stimulates scanning (*46*), eIF3i mutations impair scanning rate (*46*), while eIF3b was suggested to be involved in start-codon selection (*47, 48*). Overall, this density appears consistent in size and localization with an mRNA, indicating that the observed 43S complexes are loaded on mRNAs and may therefore be 48S complexes. As 5’UTR scanning is known to be fast (*49*), the 48S complexes we observe are likely to correspond to a later step of AUG decoding, preceding the departure of eIF2, the binding of eIF5B and the subunit joining step. This observation suggests that in cells, start codon decoding and eIF2 departure may be a limiting step of translation initiation. Mammalian 5’UTR regions are relatively long, with a median length of ∼ 175 nucleotides in mouse cells (*50*), therefore in principle possibly accommodating the simultaneous loading of several 43S complexes. Due to the high rate of the scanning step (*49*), such a situation of simultaneous loading would be expected to result in small queues of two or three 43S complexes, the first one being at the AUG codon, and the next ones accumulating afterwards. However, our subtomogram averages do not reveal close neighbors for 43S complexes suggesting that a single 43S may be loaded per mRNA, consistent with previous footprinting work (*51*).

We also observe abundant 43S complexes in 4h ANS-treated cells but the density we attribute to the mRNA is much less pronounced (Fig. S12), indicating that the loading of those 43S complexes on mRNA may be inhibited. A subclass of the 4h stress 43S complexes (551 particles) displays an extra density compatible in size and shape with eIF4G (Fig. 5E) (*44*). The 4F complex is necessary for scanning and this class may represent 43S particles that remained closer to the cap or result from an inhibition of eIF4E.

To test whether stress conditions affect the spatial distribution of the observed small subunit complexes, we visualized the different populations of 40Ss in their original tomograms and noticed that the 43S complexes seem to form small clusters especially in the 4h stress dataset (Fig. S13). Plotting the percentage of particles with 2 or more neighbors within a given distance confirmed a tendency to clump of the 43S particles in the 4h ANS dataset (and to a lesser extent in the 20min ANS dataset, Fig. S13), while the free 40S particles remain more evenly distributed. Averaging the population of 43S with 2 or more neighbors closer than 50 nm at 4h stress did not result in an increase in the density we attributed to mRNA, indicating that the clumping 43S may not be loaded on mRNA. It is unclear how 43S particles may cluster under collision stress. In conditions where translation activity is reduced, mRNA tend to clump - and ultimately form large stress granules also containing ribosomal proteins in the severe case of cells treated with initiation inhibitors (*52, 53*). Whether 43S complexes at 4h collision stress may be interacting with clumping RNA molecules in a non-canonical manner, for instance via eIF3d (*54*), remains to be further investigated.

## Conclusion

In summary, our work visualizes the spatio-temporal evolution of the translational responses to persistent collision stress in mammalian cells. Our analysis reveals that off-pathway 80S and non-functional 60S complexes accumulate in cells over time. In parallel, the shift between distinct populations of aberrant 40S complexes at different stress timepoints indicate the succession of independent initiation inhibition mechanisms. Our work highlights the value of *in situ* cryoET as a tool complementing biochemical and cellular assays for the study of translational regulation mechanisms and represents a blueprint for future *in situ* cryoET studies of translational stress responses induced upon drug treatment or in pathological conditions.

## Acknowledgments

We thank Sofia Ramalho, Joyce van Loenhout and Anjani Parag for help with preliminary experiments. We are grateful to Stuart C. Howes and Menno Bergmeijer for cryoEM support as well as to Mariska Gröllers Mulderij for support with cell culture. We thank Ramanujan Hegde and Stefan Pfeffer for insightful discussions. The work was supported by the European Research Council under the European Union’s Horizon 2020 Program (ERC Consolidator Grant Agreement 724425 - BENDER) and the Nederlandse Organisatie voor Wetenschappelijke Onderzoek (Vici 724.016.001 to FF and Veni 212.152 to JF). A.d.G was supported by the US National Institutes of Health (NIH) grant GM133598.

## Author contributions

J.F. designed the project, performed the in situ cryoET data acquisition, processing, J.F. and M.V. performed nearest neighbor analysis, J.S. performed the S35 protein synthesis experiment, S.F. performed preliminary experiments, J.F., M.V., J.S., Y.M., E.S., A.d.G., W.F. and F.F. analyzed the data and J.F. wrote the manuscript with input from all authors.

## Data and materials availability

All data are available in the main text or the supplementary materials. Electron density maps are deposited in the Electron Microscopy Data Bank with the following accession numbers: EMD-XXXXX.

## Supplementary Materials

Materials and Methods

Figs. S1 to S13

