## Supplementary Material for "Visualization of translation reorganization upon persistent collision stress in mammalian cells"

### Materials & Methods

#### Cell culture

wt MEF cells were a kind gift from M. Molinari. Cells were grown in DMEM with 10% Fetal Bovine Serum (FBS) at 37°C and 5% $CO_2$. Cells were treated with 200nM anisomycin for 15min, 1h or 4h prior to experiments.

#### Western blotting antibodies

The cell surface was briefly washed with PBS and the cells were harvested in trypsin (+ANS for stressed conditions). Cells were washed once in ice cold PBS and pellets were snap frozen until further use. Cell lysates were prepared by solubilization of the cell pellet in TEN Triton buffer (20mM Tris pH 7.5, 150mM NaCla, 1mM EDTA), supplemented with Protease Inhibitor (Roche) and phosphatase inhibitor (Cell Signaling #5870), at 4°C for 30min. Cell lysates were centrifuged at 20 000g for 20 min and the supernatant was diluted in 2X sample DTT-loading buffer. Samples were heated to 95 degrees for 5min and stored at -20. For western blotting we used following commercial antibodies: eIF2α (Cell Signaling Antibody #9722), phospho-eIF2α (Cell Signaling #9721), phospho-p38 (Cell Signaling #4511), phospho-JNK1+2 (Cell Signaling mAb #4668), caspase 3 (Cell Signaling Antibody #9542), PARP-1 (Cell Signaling Antibody #9662), GCN1L1 (Thermo Fisher Scientific A301-843A-M), EDF1 (Abcam #ab174651), ZNF598 (Thermo Fisher Scientific A305-108A-M), RPL35 (Life Technologies PA552245), RPS10 (Abcam #ab151550). Relative band intensity quantification was performed in Fiji/ImageJ, their background was subtracted and

#### S35 protein synthesis

MEF cells were grown in DMEM FBS and treated with 100$\mu$g/mL CHX or 200nM ANS for 15min, 1h, or 4h prior to labelling. Cells were then treated with DMEM methionine-free media (ThermoFisher Scientific #21013024) for 20 min and incubated with 30 µCi/ml 35S-methionine label (Hartmann Analytic) for 1 h. After washing the samples with PBS, proteins were extracted with lysis buffer (50 mM TrisHCl pH 7.5, 150 mM NaCl, 1% Tween-20, 0.5% NP-40, 1× protease inhibitor cocktail (Roche) and phosphatase inhibitor cocktail (Sigma Aldrich) and precipitated onto filter paper (Whatmann) with 25% trichloroacetic acid and washed twice with 70% ethanol and twice with acetone. Scintillation was then read using a liquid scintillation counter (Perkin Elmer) and the activity was normalized by total protein content. All experiments were done in technical triplicates for each biological unit.

#### Polysome profiling

MEF cells were grown to about 70% confluency and harvested with trypsin (+ANS in the stress conditions). Cells were washed once in ice cold PBS (+ANS) prior to lysis for 30min on a rotated wheel at 4°C in an adapted RIPA buffer (50mM Hepes KOH pH7.4, 15mM MgOAc, 100mM KOAc, 5% glycerol, 0.1% triton X-100, 0.1% SDS, 0.5% sodium deoxycholate, 1mM DTT, 1mM PMSF, protease inhibitor tablet). Lysate was cleared by centrifugation at 8000 g for 5 min and added on top of a 10-50% sucrose gradient (in buffer: 25mM Hepes-KOH, 100mM KOAc, 5mM MgOAc, 1mM DTT, 0.5mg/mL heparin, 1mM PMSF). Gradients were centrifuged for 3h at 32 000 rpm in a 32.1 Ti rotor (Beckman). Gradients were sampled and OD was measured using a gradient station (Biocomp).

Grid preparation

MEF were seeded on R2/2 holey carbon on gold grids (Quantifoil or Protochips) coated with fibronectin in a glass bottom dish (Mattek or Ibidi) and incubated for ~24 h. For stress conditions, cells were then incubated with 200 nM anisomycin in DMEM + 10% FBS for 20min, 1h or 4h. Grids were incubated for 5min in DMEM (+ 200 nM ANS for stressed conditions) with 10% dextran-40 used as non cell permeable cryo-protectant (*55*). Grids were then immediately mounted to a manual plunger, blotted from the back for ∼10 s and plunged into liquid ethane.

Lamella preparation

Lamellae were prepared using an Aquilos FIB-SEM system (Thermo Fisher Scientific). Grids were sputtered with an initial platinum coat (10 s) followed by a 10 s gas injection system (GIS) to add an extra protective layer of organometallic platinum. Samples were tilted to an angle of 15° to 22° and 10 µm wide lamellae were prepared. The milling process was performed with an ion beam of 30 kV energy in 3 steps: 1) 500 pA, gap 3 µm with expansion joints, 2) 300 pA, gap 1 µm, 3) 100 pA, gap 500 nm. Lamellae were finally polished at 30-50 pA with a gap of 200 nm.

Data collection for non-treated, 20min and 1h ANS datasets

A total of respectively 93, 48 and 74 tilt series were acquired on a Talos Arctica (Thermo Fisher Scientific) operated at an acceleration voltage of 200 kV and equipped with a K2 summit direct electron detector and 20eV slit energy filter (Gatan). Images were recorded in movies of 5-8 frames at a target defocus of 4 to 6 µm and an object pixel size of 2.17 Å. Tilt series were acquired in SerialEM using a grouped dose-symmetric tilt scheme covering a range of ±54° with a pre tilt of ±10° and an angular increment of 3°. The cumulative dose of a series did not exceed 80 e-/Å2.

Data collection for 4h ANS dataset

A total of 53 tilt series were acquired on a Titan Krios (Thermo Fisher Scientific) equipped with a K3 summit direct electron detector and Bioquantum energy filter (Gatan). The microscope was operated at an acceleration voltage of 300 kV and 20eV slit. Images were recorded in movies of 10 frames at a target defocus of 4 to 6 µm and an object pixel size of 2.17 Å. Tilt series were acquired in SerialEM (*56*) using a grouped dose-symmetric tilt scheme covering a range of ±54° with a pre tilt of ±10° and an angular increment of 3° (*57*). The cumulative dose of a series did not exceed 80 e-/Å2.

Tomogram reconstruction

Movie files of individual projection images were motion- and CTF-corrected in Warp and combined into stacks of tilt series (*58*). The combined stacks were aligned using patch tracking in IMOD (*59*). CTF estimation for entire tilt series was performed in Warp (*58*) and full tomograms were reconstructed by weighted back projection at a pixel size of 17.36 Å. Ice thickness was determined manually and was found to be <200nm for all lamellae.

#### Particle picking

Particle coordinates were determined using PyTom (*60*) template matching against a reconstruction of a mammalian 80S, 40S or 60S ribosomes, filtered to 40 Å. The determined positions of ribosomes were used to extract subtomograms and corresponding CTF volumes at a pixel size of 8.68 Å (4× binned) in Warp (*58*) for the 80S ribosomes and 4.34 Å (2× binned) for the 40S and 60S particles.

#### Subtomogram analysis

The extracted subtomograms were used for 3D classification with image alignment against a low pass filtered 80S/60S/40S ribosome map as reference in RELION (3.1.4) (*61*) to exclude false positives. The remaining ribosome subtomograms were refined in RELION and good 80S particles were re-extracted in Warp at a pixel size of 4.34 Å (2x binned). Bin2 80S subtomograms were refined in RELION with a mask on the LSU prior to a first round of 3D classification without image alignment with a mask on the SSU to separate rotated from non-rotated ribosomes. A second round of classification was performed using a mask positioned on the tRNA and elongation factors sites, optimizing the mask extension and class number to yield stable classes. The classes containing a high number of particles (decoding E) were further submitted to 3D classification without alignment with 6-10 classes to analyze the presence of smaller sub-populations (like POST and POSTi). The different classes were finally subjected to an iterative refinement in M (*62*).

Bin2 40S and 60S subtomograms were submitted to 3D refinement in RELION followed by rounds of 3D classification without image alignment, introducing masks on densities visible at low threshold, in order to separate the corresponding particles.

#### Spatial analysis

For each 80S ribosome particle, the position and angles from RELION star file was used to calculate the position of the mRNA entry and exit sites and distances were calculated from its mRNA entry site to all other particles mRNA exit sites; the shortest distance was retained. Conversely, the shortest distance of all other entry sites to a particle exit site was also computed. The distribution of these shortest distances was plotted using matplotlib.

For 43S clustering analysis, a similar approach was used relying only on the position of the particle center of mass as in relion starfiles.

#### Visualization

Membranes were segmented using Tomosegmemtv (*63*). All figures were prepared in UCSF ChimeraX (*64*) using the ArtiaX pluggin (*65*) for mapping back subtomogram averages to their coordinates in the original tomogram.

### Supplementary Figures

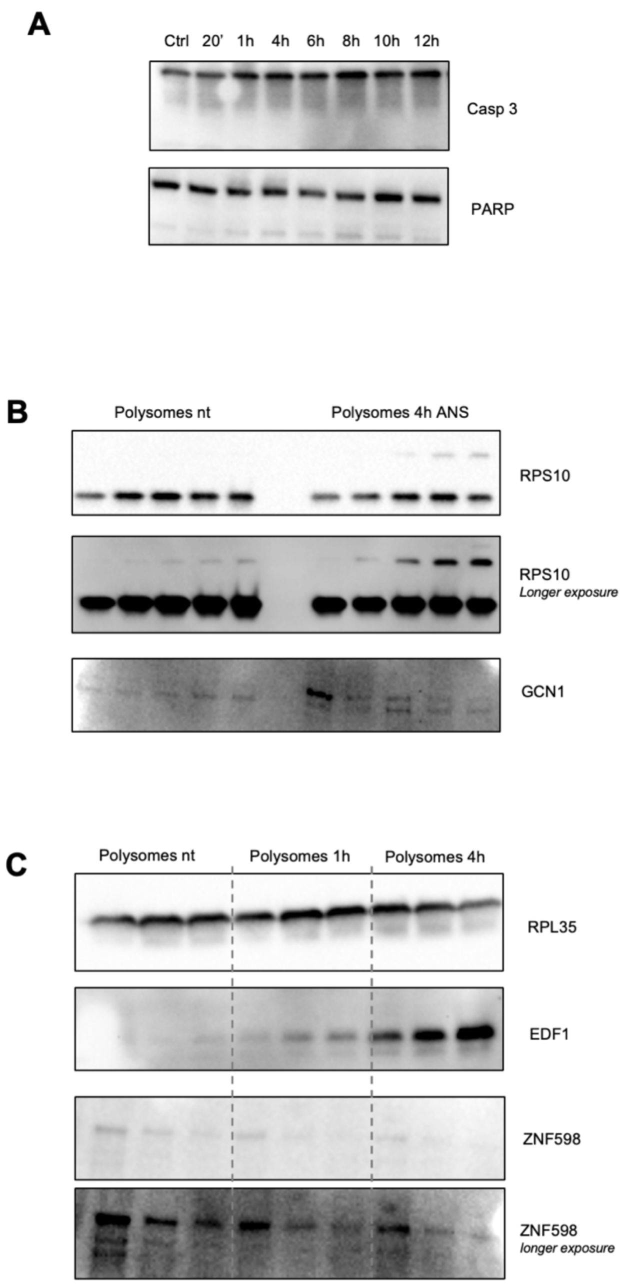


**Supplementary Figure S1: Western blot analysis**. (A) Analysis of the possible cleavage of apoptosis markers casp3 and PARP followed over time of low dose ANS stress. (B) Analysis of the presence of known ribosome collision sensors on polysomes of control (nt) cells and cells stressed with low dose ANS for 1h and 4h. RPL35 is used as loading control antibody, the presence of EDF1 and ZNF598 is tested on polysomes. (C) Detection of RPS10 ubiquitination and GCN1 presence on polysomes of nt and 4h stressed cells.

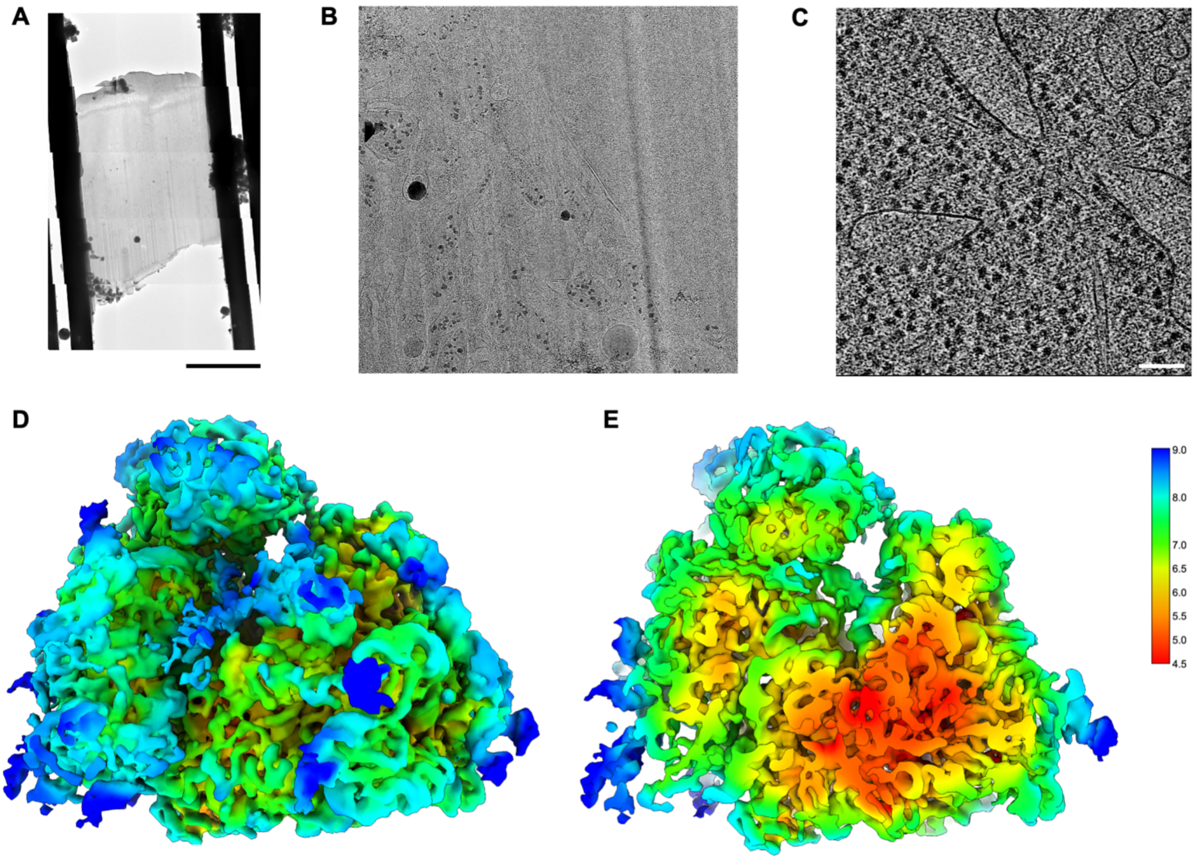


**Supplementary Figure S2: In situ FIB-CET workflow**. (A) TEM map of a lamella through a MEF cell. Scale bar 5 $\mu$m. (B) Close up view on the lamella map displaying cellular features. (C) Slice through a representative tomogram, scale bar 100 nm. (D) 80S ribosome subtomogram average colored according to local resolution. (E) Same map as in D, clipped for visualization of the internal best resolved part of the LSU. Average resolution is 5.9 Å and locally down to 4.5 Å.

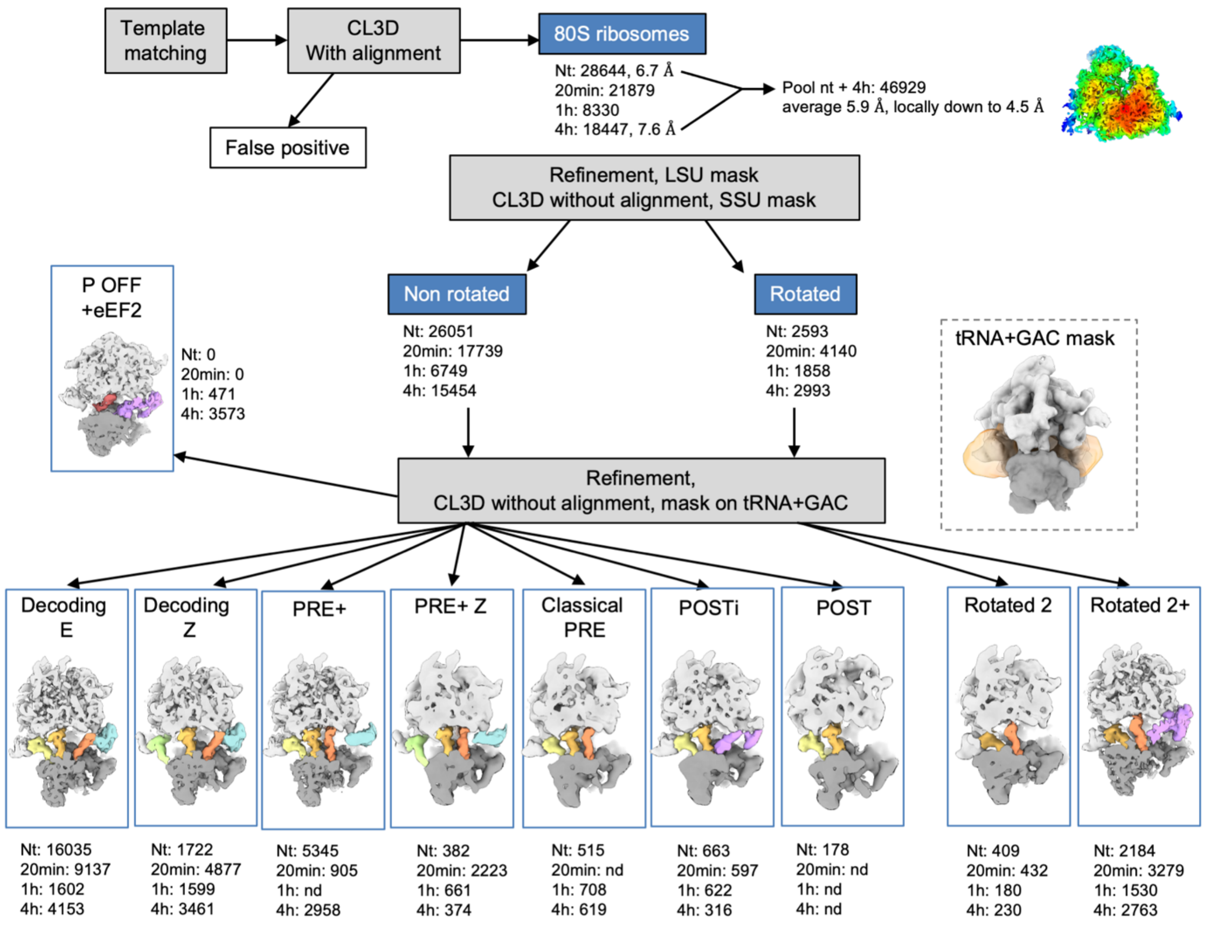


**Supplementary Figure S3: Cryo-ET data analysis workflow.** Particles were picked using template matching in PyTom (*60*), subtomograms extracted in Warp (*58*) and false positive were eliminated using CL3D with image alignment in RELION (*61*), followed by subtomogram alignment with a mask on the LSU. The particles were hierarchically classified with CL3D without image alignment, first using a mask on the SSU to separate rotated from unrotated states, and then using a mask on the tRNA sites and the GAC. Final classes were refined in M (*62*).

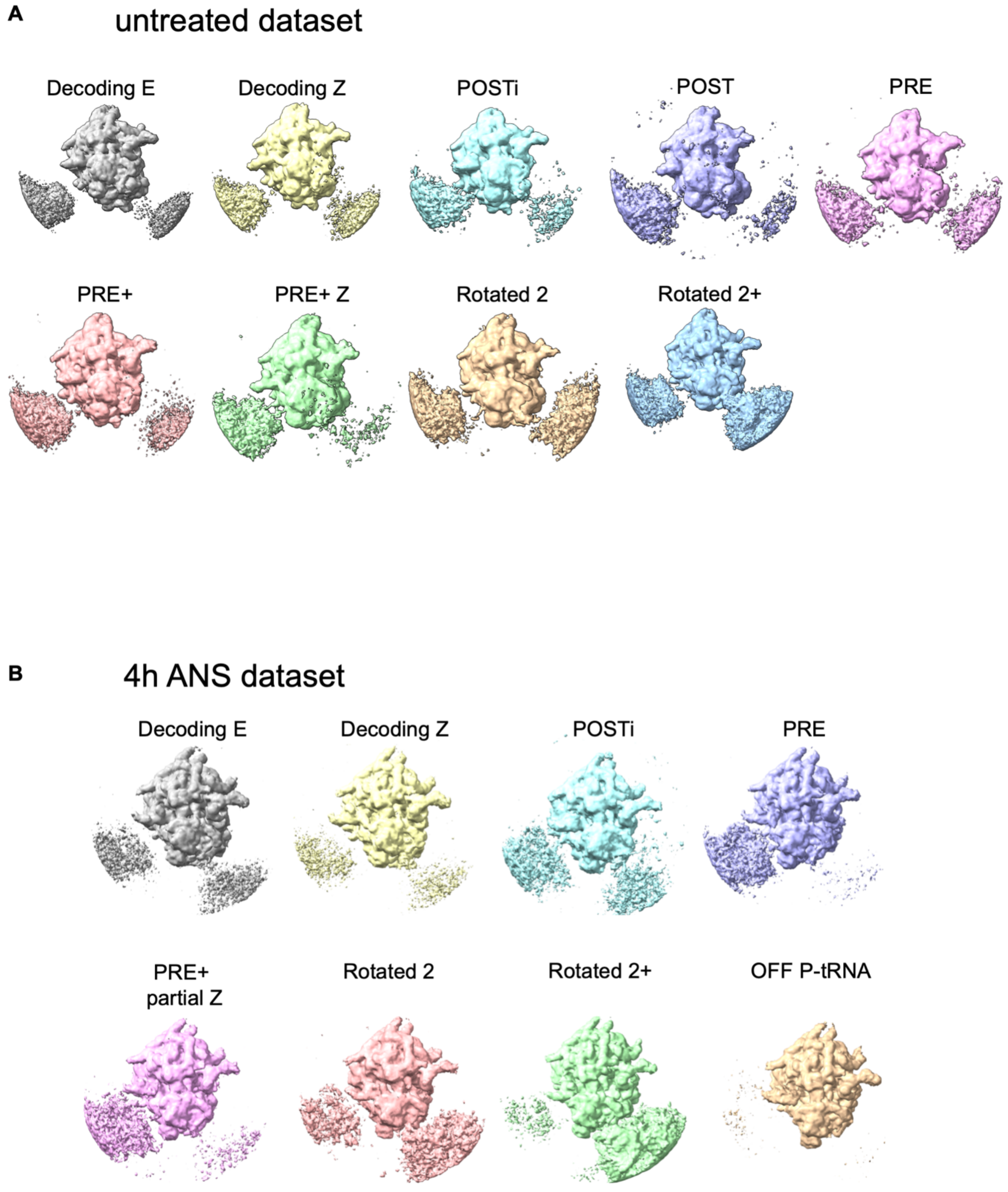


**Supplementary Figure S4: Overview of leading and trailing neighbor densities in the main 80S classes.** (A) in the untreated dataset, (B) in the 4h ANS dataset. Unmasked maps from relion Refine3D jobs are displayed.

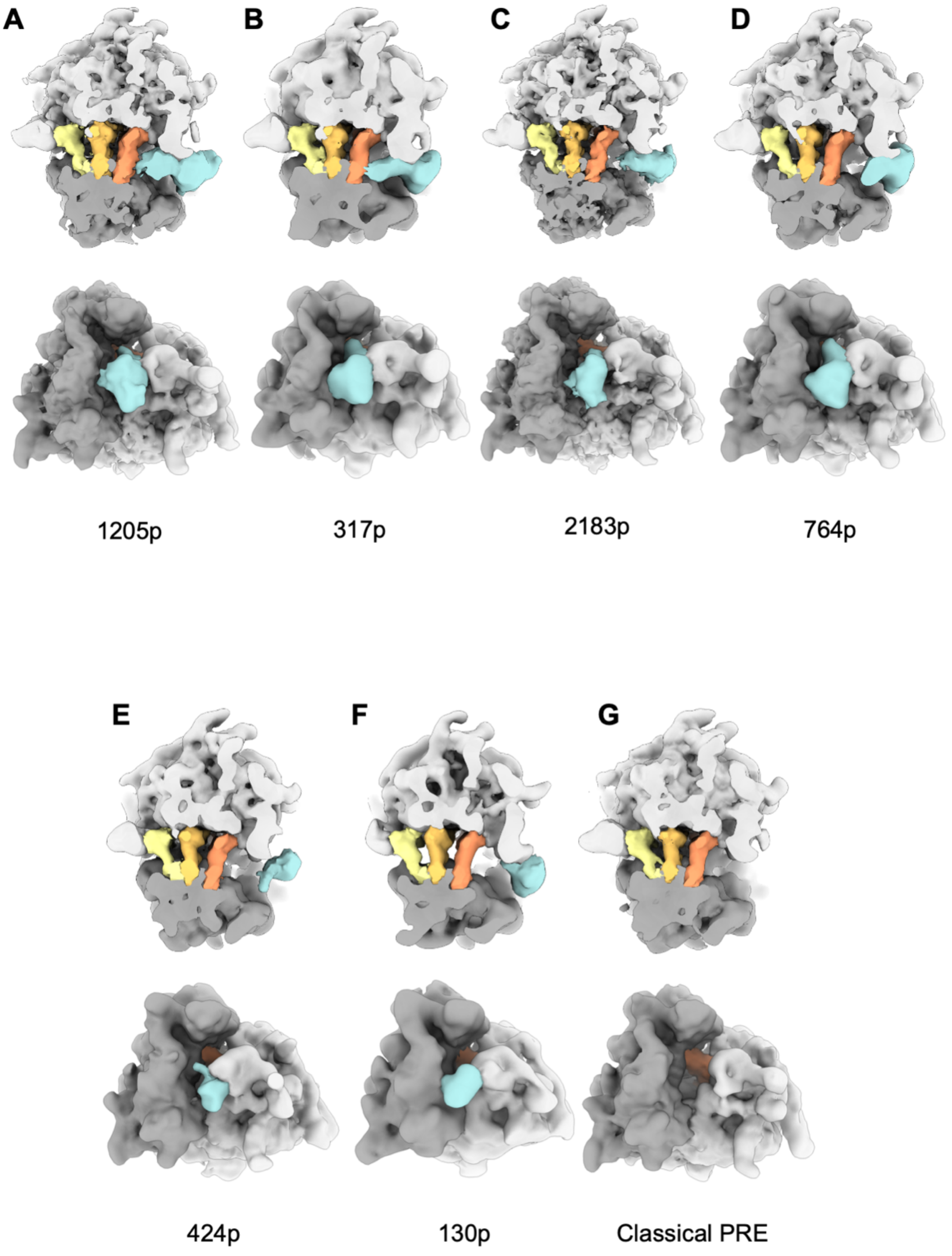


**Supplementary Figure S5: Sub-classes observed of the PRE+ state**. The ribosome is clipped as in Fig 2C. and the same color code is used. The number of particles in each class is indicated below the corresponding subtomogram average.

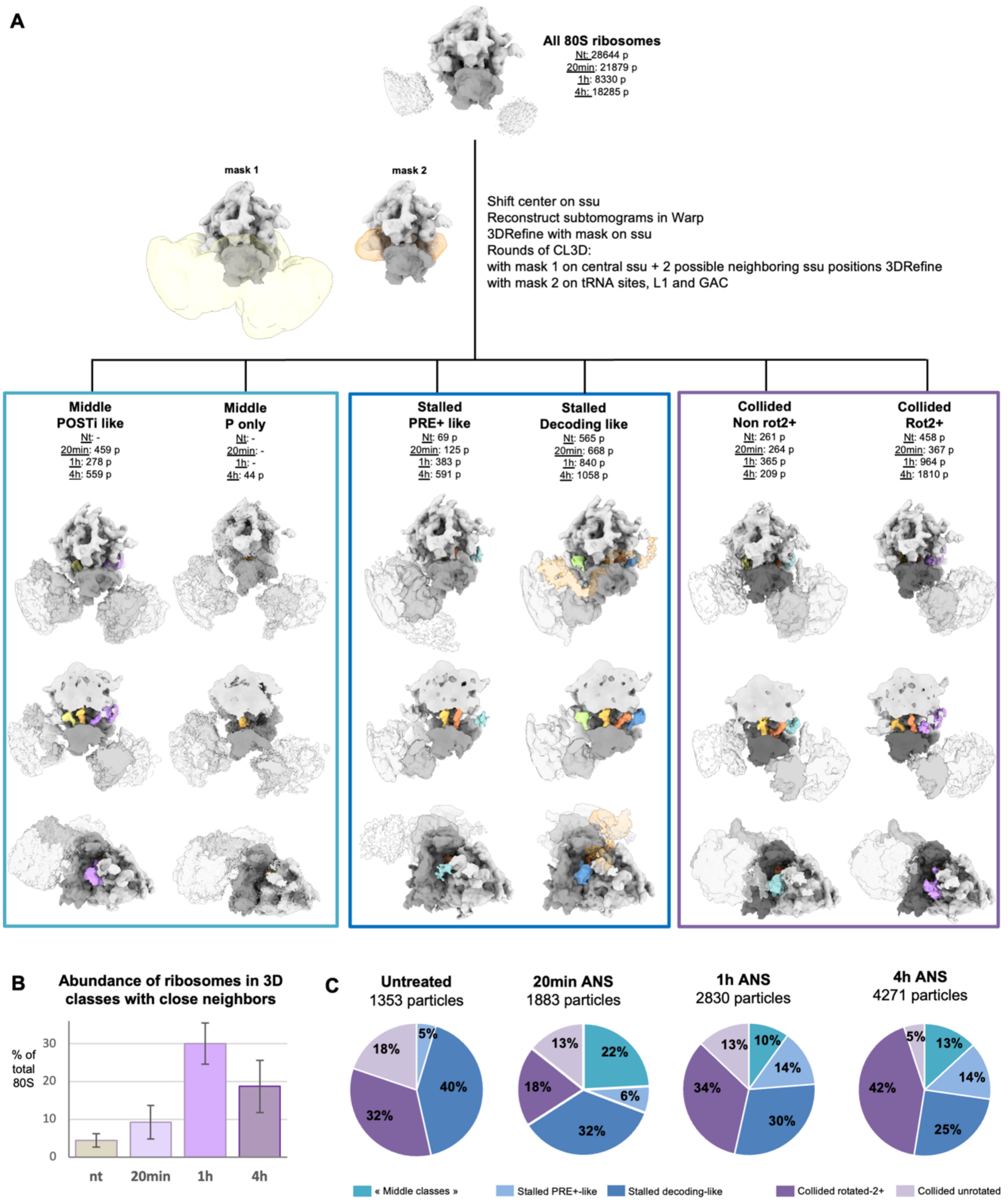


**Supplementary Figure S6: Sub-classes observed for the ribosomes with close neighbors**. (A) 3D classification workflow and final classes subtomogram averages. The classes with 2 close neighbors appearing as slightly less defined densities are boxed in cyan. The classes corresponding to typical leading ribosomes in a disome are boxed in dark blue and the ones corresponding to typical collided ribosomes are boxed in dark purple. The same class is shown in 3 different views in column: a top view, the same view clipped for visualization of the internal tRNA, and a side view to visualize the GAC. (B) Rough quantification of the abundance of ribosomes with close neighbors as estimated by 3D classification in RELION. Bar and whiskers are mean and s.d. across tomograms (untreated n = 87, 20 min ANS n = 68, 1h ANS n = 36, 4h ANS n = 45). (C) Relative abundances of the different classes visualized in (A).

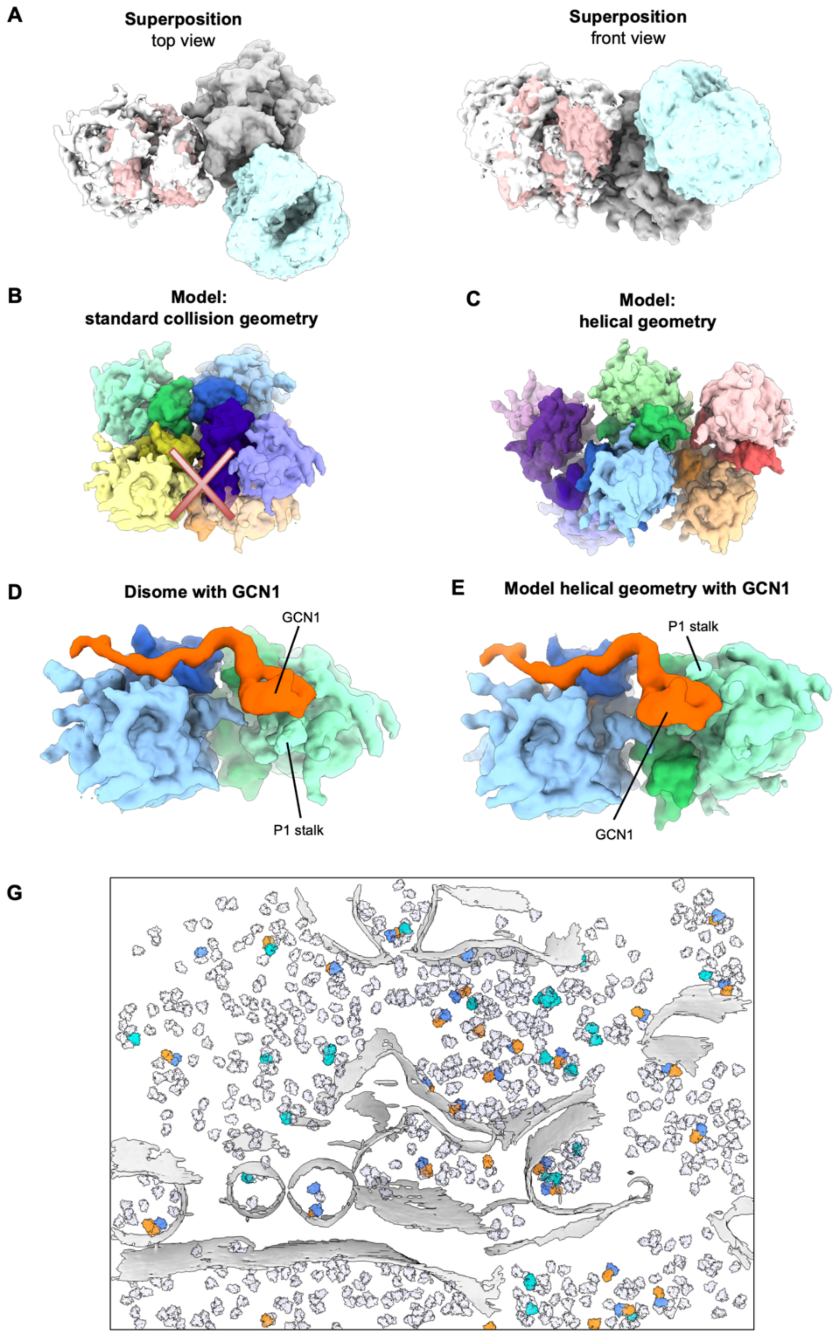


**Supplementary Figure S7: Comparison of the distinct classes of ribosomes with close neighbors**. (A) Superposition of the subtomogram averages of ribosomes with a collided neighbor (in white) or 2 helical neighbors in cyan and pink. (B) Model of a chain of 5 ribosomes in the collision geometry, displaying clashes for n ≥ 4. (C) Model of a chain of 7 ribosomes following the helical geometry, no clash. (D) Typical collided disome with GCN1 bound. (E) Model of a disome with helical geometry and GCN1 bound to the leading ribosome, revealing incompatibility of this geometry with GCN1 simultaneously binding to the trailing ribosome. (G) GCN1 bound stalled (blue) and collided (orange) ribosomes, as well as helical geometry ribosomes mapped back onto their original tomograms. Segmented membranes are displayed in light gray. Other 80S ribosomes are displayed in transparent light mauve.

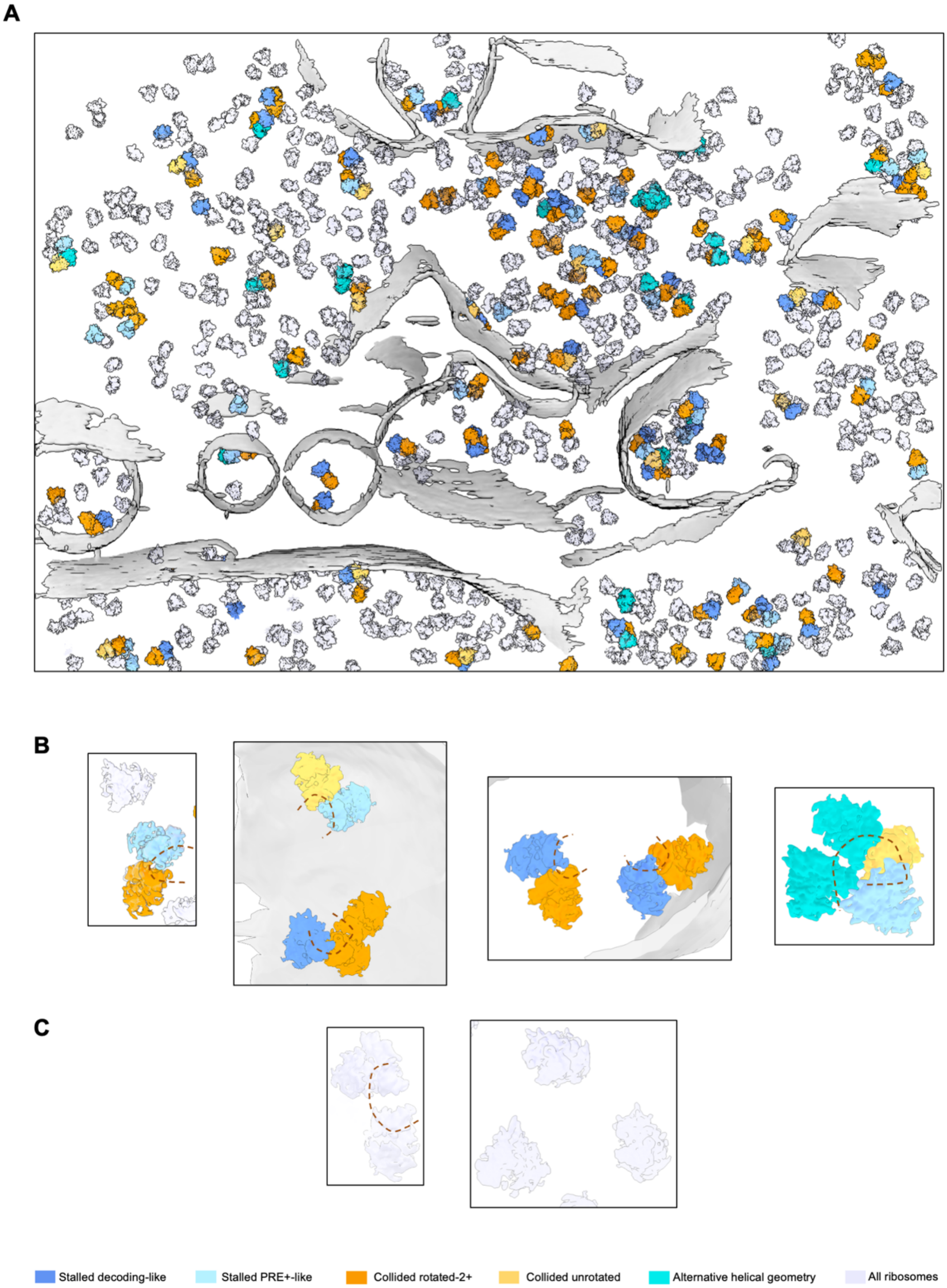


**Supplementary Figure S8: Views of ribosomes collisions**. (A) All collided and helical ribosomes mapped back to tomogram, colored according to their class. (B) Close up views of cytosolic and ER-bound ribosome collisions. (C) Close up views of 80S with and without polysome neighbors.

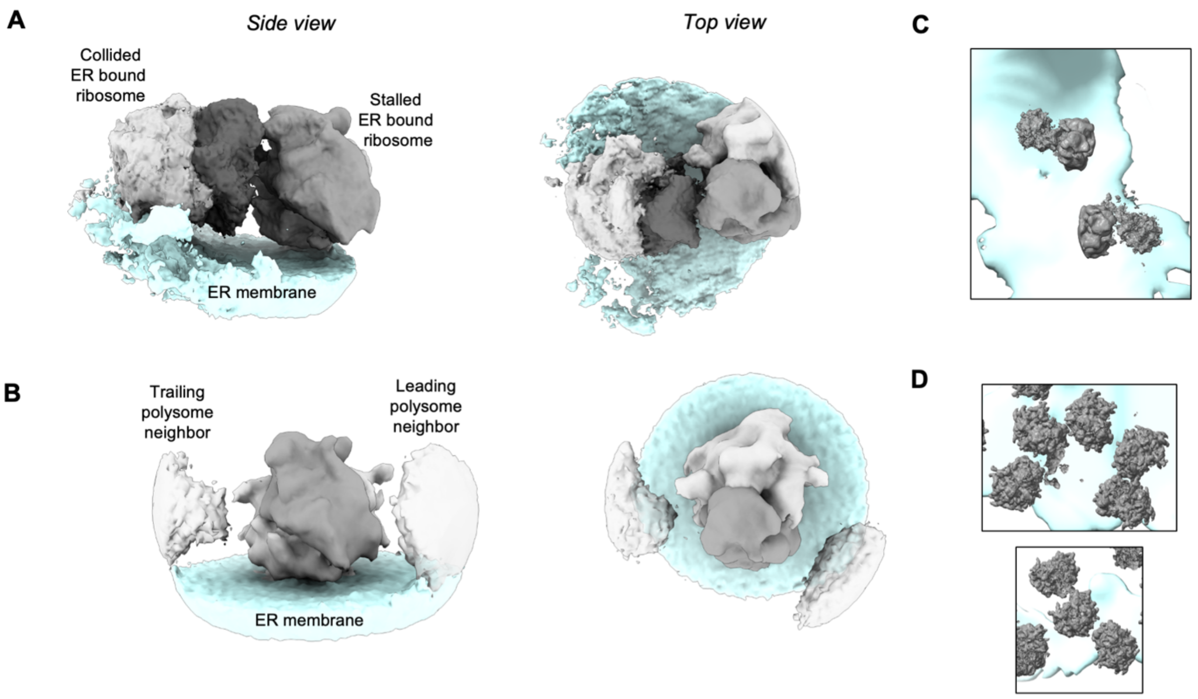


**Supplementary Figure S9: Views of ER-bound ribosomes collisions**. (A) Subtomogram average of ER-bound collisions. ER membrane is depicted in light cyan. (B) Close-up view of mapped back ER-bound collided disomes.

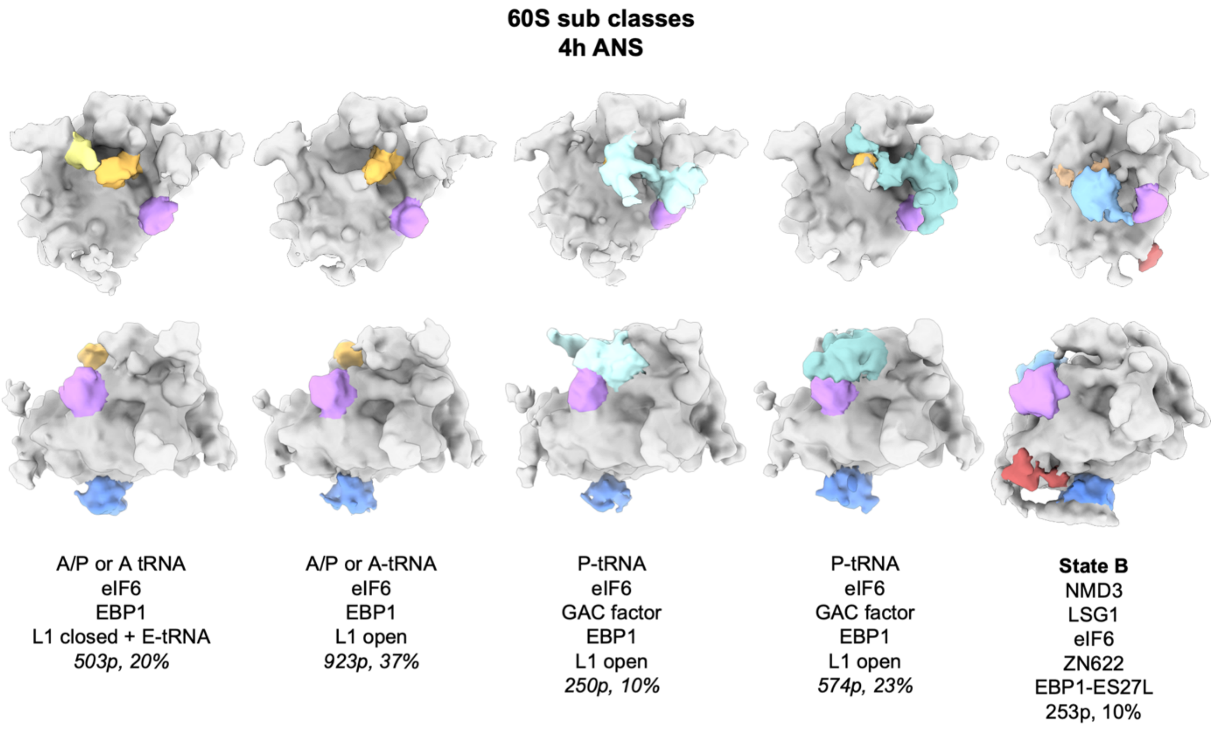


**Supplementary Figure S10: 3D classes of 60S complexes in the 4h low dose ANS stress dataset.** 2 views are shown in column for each class: a top view and a side view. Presence of tRNAs and extra factors bound is indicated, as well as the observed position of the L1 stalk, the amount of particles in each class and its relative abundance.

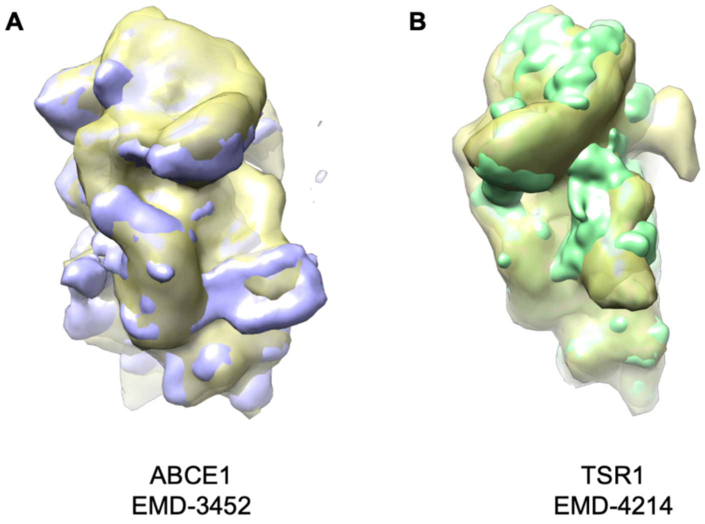


**Supplementary Figure S11: Analysis of the 4h ANS 40S complex with extra eIF5B-like density.** (A) Superposition of the complex observed in this work (transparent yellow) with a low pass filtered ABCE1 containing complex (EMD-3452, purple). (B) Superposition with a low pass filtered TSR1 containing complex (EMD-4214, green).

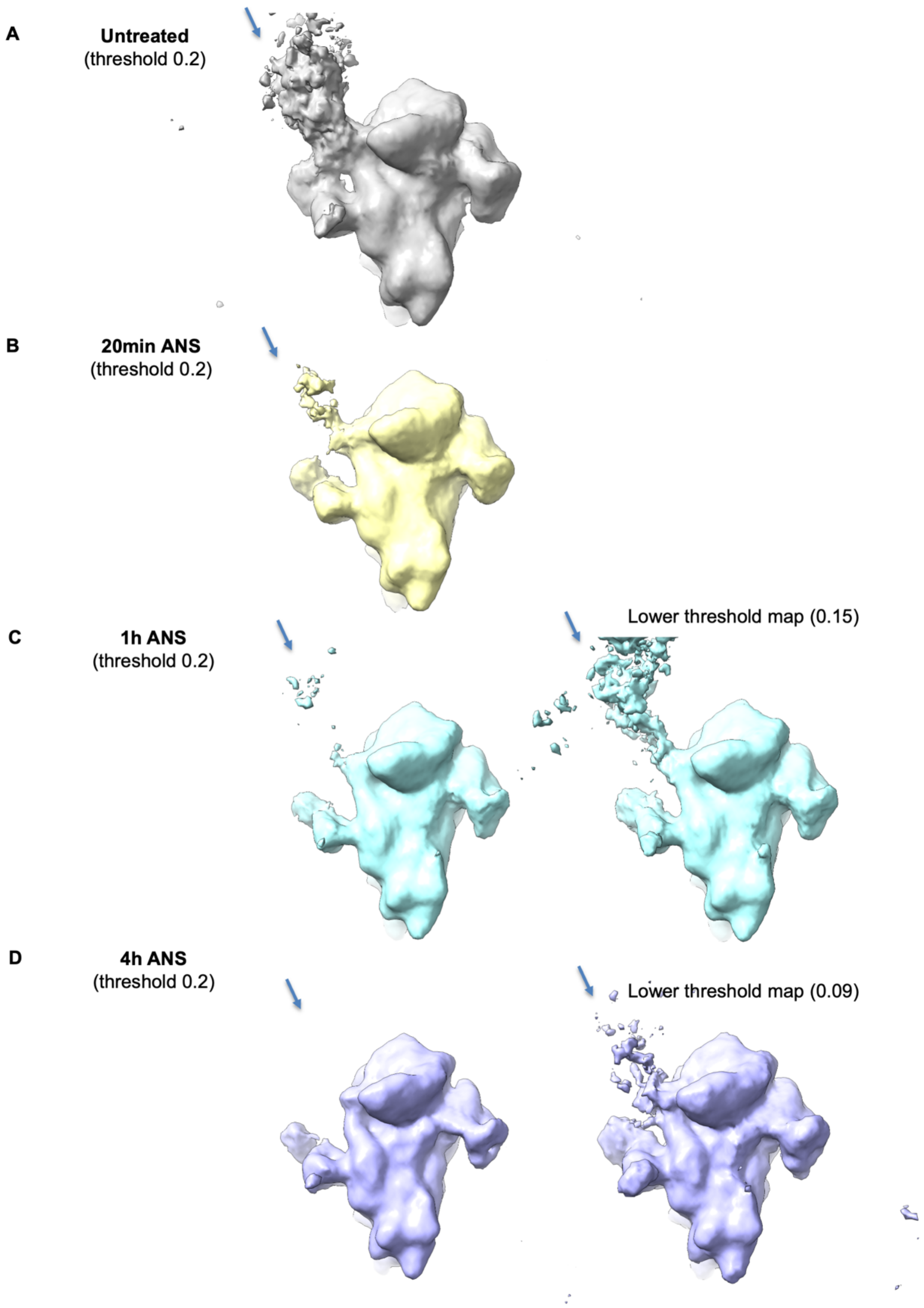


**Supplementary Figure S12: Analysis of mRNA density on 43S complexes.** Subtomogram average of 43S complexes in (A) untreated cells, (B) 20 min ANS, (C) 1h ANS, (D) 4h ANS.

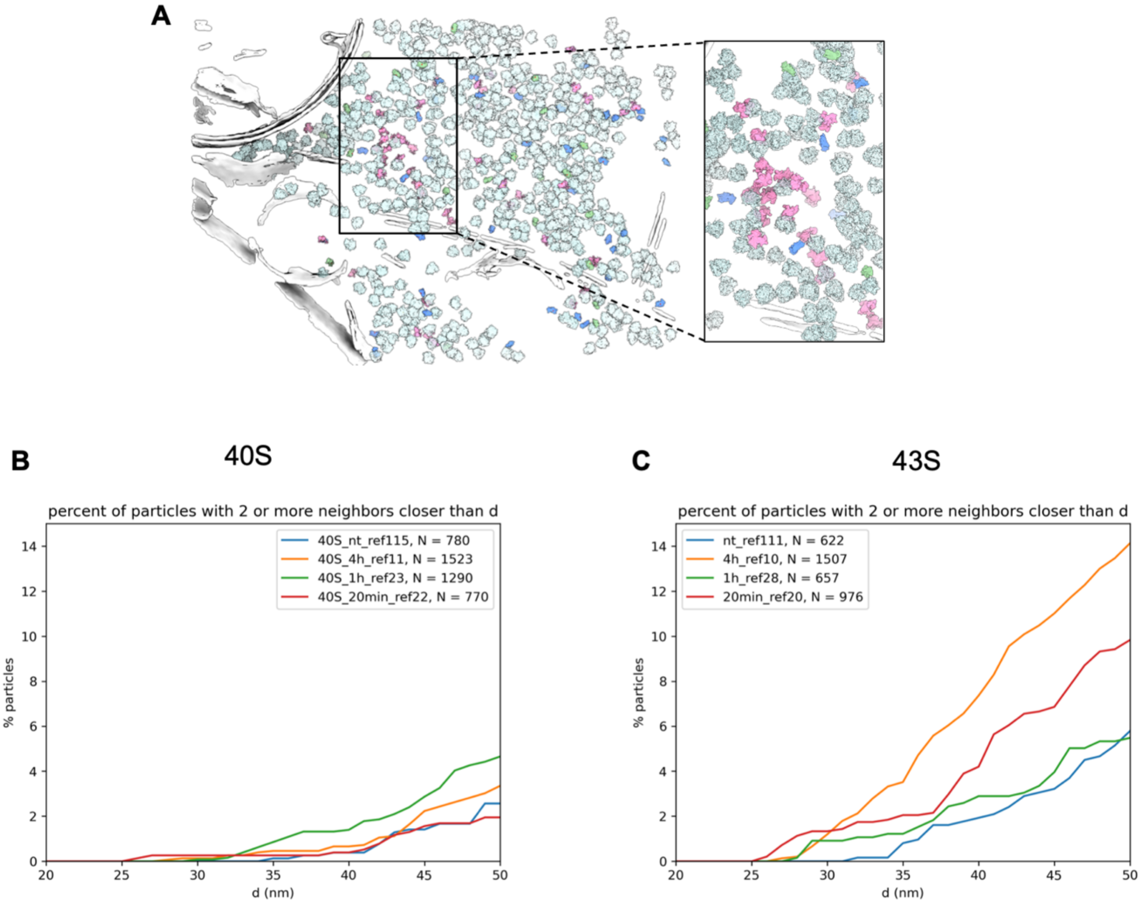


**Supplementary Figure S13: Analysis of 40S and 43S spatial distribution**. (A) 40S (blue and green) and 43S (pink) complexes mapped back onto a tomogram at 4h low dose ANS stress. 80S ribosomes are displayed in transparent light cyan and segmented membranes and microtubules are depicted in white. Close-up view on a 43S cluster. (B) Plot of the proportion of 40S particles (in %, y axis) with 2 or more neighbors closer than a distance d (in nm, x axis) across all datasets, (C) Same plots for 43S particles for all datasets.
